# The Relative Contribution of Color and Material in Object Selection

**DOI:** 10.1101/393470

**Authors:** Ana Radonjić, Nicolas P. Cottaris, David H. Brainard

## Abstract

Information about color, material, texture and shape guide how we interact with objects. We developed a paradigm that quantifies how two object properties (color and material) combine in object selection. On each experimental trial, observers viewed three blob-shaped objects — the target and two tests — and selected the test that was more similar to the target. Across trials the target was fixed while the tests varied in color and material. We present a novel observer model that allows us to describe observers’ selection data in terms of (1) the underlying perceptual stimulus representation and (2) a color-material weight, which quantifies the relative importance of color vs. material in selection. We document large individual differences in the color-material weight. Furthermore, our analyses reveal limits on how precisely selection data simultaneously constrain perceptual representations and the color-material weight. These limits should guide future efforts towards understanding the multidimensional nature of object perception.

## Introduction

In daily life, we rely on vision to select objects for a variety of goal-directed actions. For example, when we crave strawberries, we use color to decide which berries are the ripest (Figure 1A); when we sip coffee, we use glossiness to judge whether a cup is made of porcelain or paper, which in turn affects how we handle it (Figure 1B). Indeed, we continually use visual information to effortlessly and confidently judge object characteristics. Instances in which vision misleads us are sufficiently rare to be memorable, as in the case of a deflated basketball sculpture made of glass (Figure 1C).

**Figure 1.**
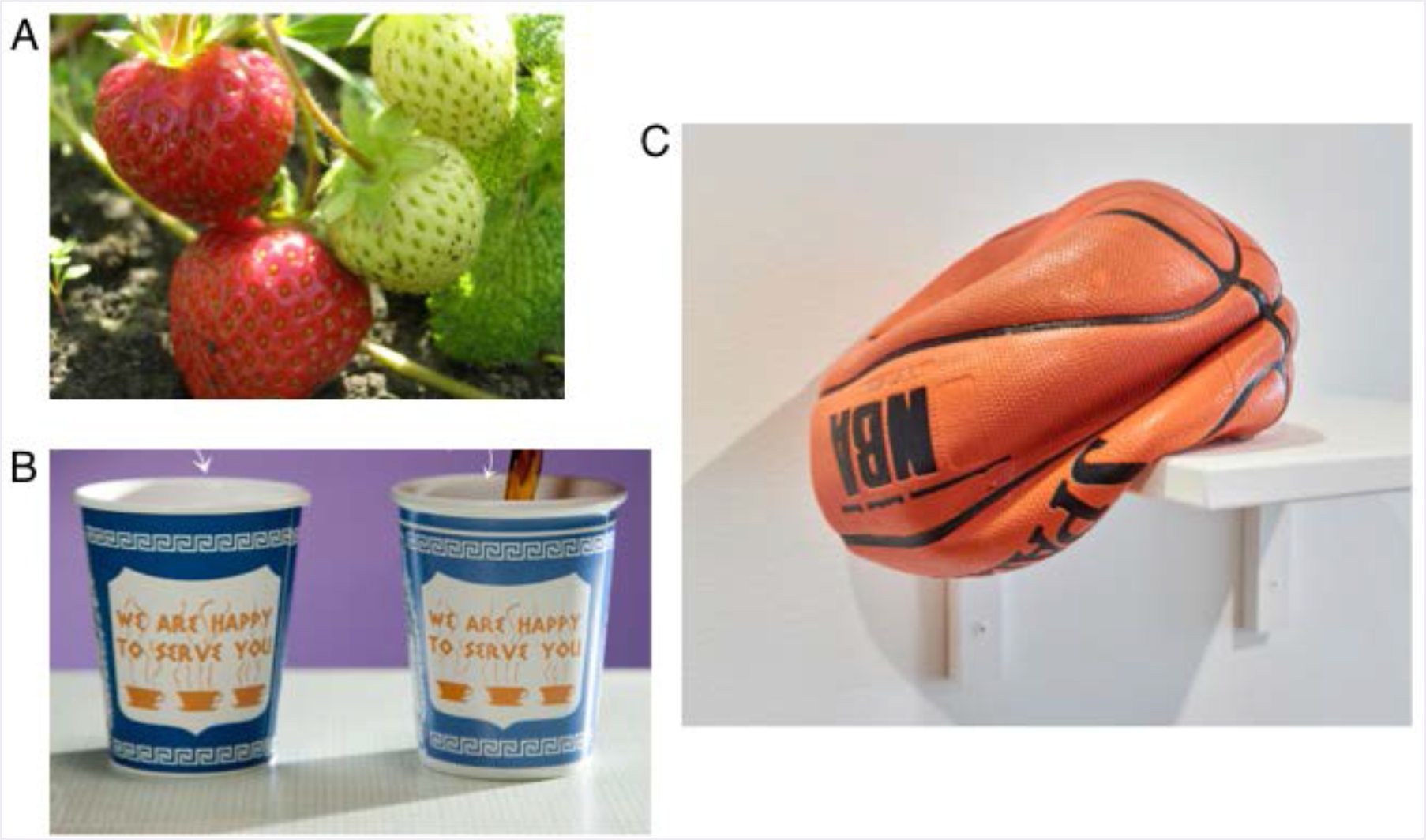
Vision is used to judge object properties. A. Color differentiates between ripe and unripe strawberries. B. Glossiness differentiates between paper and ceramic cups. C. A carefully crafted case where vision misleads us about material. What looks like a rubber basketball is in fact a sculpture made of glass, created by Christopher Taylor. (Photography sources: A: https://www.deviantart.com/regretsofblood, B: https://www.vat19.com, C: http://www.craftadvisory.com; we are currently in the process of obtaining permissions for using these photographs).

Extracting information about object properties from the image formed on the retina by light reflected from objects is a challenging computational problem. This is because the process of image formation entangles information about the intrinsic properties of objects (such as color or material) with information about the conditions under which they are viewed. For example, the retinal image is affected by changes in the illumination, the objects’ position and pose, and the viewpoint of the observer. Understanding the perceptual computations that transform the retinal image into stable representations of objects and their properties is a longstanding goal of vision science.

A large literature has employed a “divide and conquer” strategy to investigate the perception of object properties: different object attributes (color, texture, material, shape, etc.) have each been studied within their own subfields. This approach has leveraged well-controlled laboratory stimuli and relatively simple psychophysical tasks to build a quantitative understanding of how information is transduced and represented early in the visual pathways (Wandell, 1995; Rodieck, 1998). In addition, careful case studies have provided insight into how stable perception of object properties may be achieved when the experimental conditions are relatively simple and well-specified (Olkkonen & Ekroll, 2016; Foster, 2011; Chadwick & Kentridge, 2015; Fleming, 2014; Koenderink & van Doorn, 2004). Our work builds on the foundations provided by this approach and aims to extend the study of object perception in two critical ways. First, we want to move beyond highly-simplified laboratory stimuli and tasks and devise paradigms in which object perception is probed using both naturalistic stimuli and naturalistic tasks. Second, to explain real-life object representations, we need to describe how perceptual judgments along multiple dimensions (shape, color, material, texture, etc) combine and interact.

Our experimental paradigm employs naturalistic stimuli in combination with a two-alternative forced-choice object selection task. This task captures a core aspect of how vision is used in real life, where it guides object selection in service of specific goals (e.g., selecting nutritious and avoiding spoiled food Radonjić, Cottaris, & Brainard, 2015b). We have previously shown how a version of the elemental selection task can be embedded within more complex and naturalistic tasks to probe color perception (Radonjić, Cottaris, & Brainard, 2015a; Brainard, Cottaris, & Radonjić, 2018). Here we elaborate the selection task to measure the underlying perceptual representations of both object color and material (specifically, glossiness) and quantify how these two perceptual dimensions combine in object selection.^1^

**Figure 2.**
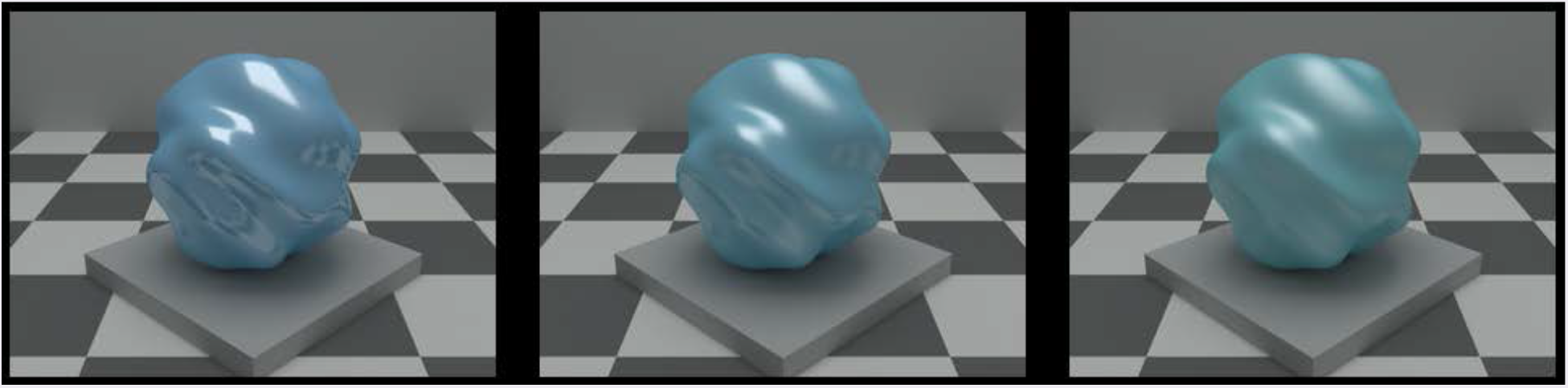
Example trial in the experiment. The observer viewed three rendered scenes, each containing a blob-shaped object. The object in the center scene was the target object. The objects in the left and the right scenes were the test objects. The observer selected which of the two test objects was most similar to the target. The two tests differed from the target in color and/or material (see text for details). In the example shown, both tests differ from the target in both color and material (left test, C+3M_3, is more glossy and bluer; right test C-3M+3 is more matte and greener; see Figure 3 for explanation of stimulus coding conventions).

The object selection task is illustrated by Figure 2. On each trial, observers viewed three blob-shaped objects — the target and two tests — and selected the test that was more similar to the target. Across trials, the color and glossiness of the target object was fixed. The tests were identical to the target in their shape, size and pose, but their color and material varied from trial to trial. Test color could vary from blue to green, across 7 different levels; test material could vary from matte to shiny, also across 7 levels (Figure 3). For each pair of test objects, we measured the probability that each member of the pair was selected.

**Figure 3.**
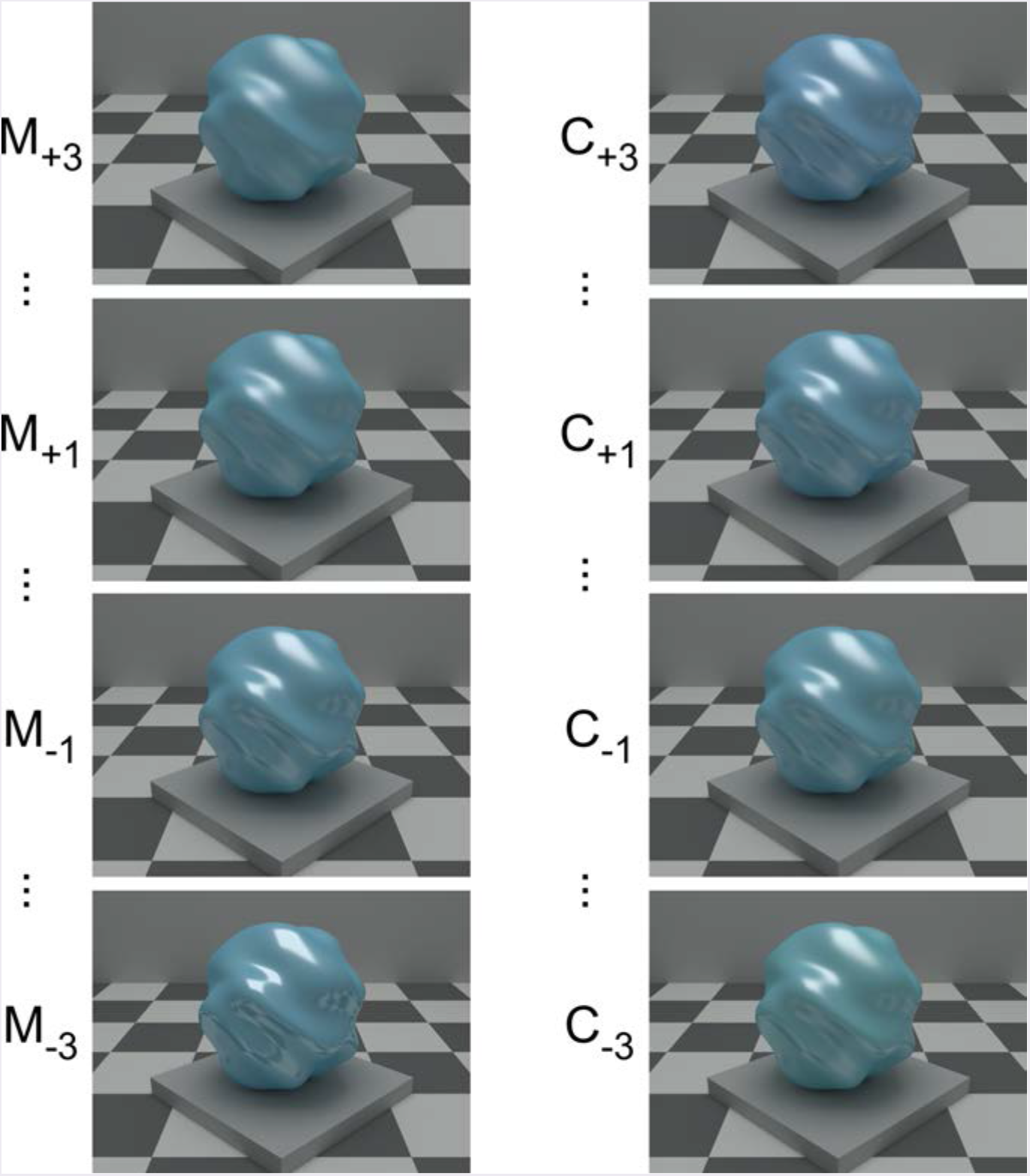
Stimulus examples. Left column shows stimuli that have the same color as the target and vary only in material: these illustrate variation on the material dimension, from most glossy (M_-3_) to most matte (M_+3_). The numerical labels indicate the nominal degree of color/material difference from the target, which is labeled C_0_M_0_. Only large (M_-3_, M_+3_) and small (M_-1_, M_+1_) material difference steps are shown. Right column shows stimuli that have the same material as the target and vary only in color: these illustrate variation on the color dimension, from greenest (C_-3_) to bluest (C_+3_). Only large (C_-3_, C_+3_) and small (C_-1_, C_+1_) color difference steps are shown.

We report three primary results. The first is theoretical. To understand the selection data, we need an observer model that translates the raw data into an interpretable form. An important advance of the work we present here is the development of such a model. The model describes the data in terms of how the stimuli are positioned along underlying perceptual dimensions and how distances along these dimensions are combined to guide object selection. Our second result is experimental. We show that there are large individual differences in the degree to which observers rely on object color relative to object material in selection. Some observers base their selections almost entirely on color, some weight color and material nearly equally, and others rely almost entirely on material. Third, a fine-grained analysis of our data, in parallel with model comparisons, clarifies limits on how precisely selection data may be leveraged to simultaneously reason about perceptual representations and color-material trade-off. These limits, which we make explicit, are important to recognize as we and others move towards understanding the multi-dimensional nature of object perception. Below, we present each of these results in detail.

## Results

### Perceptual model

Our observer model builds directly on our recent work on color selection (Radonjić, Cottaris, & Brainard, 2015b; Radonjić, Cottaris, & Brainard, 2015a; Brainard, Cottaris, & Radonjić, 2018) and incorporates concepts from multidimensional scaling (Kruskal, 1964), the theory of signal detection (Green & Swets, 1966), and maximum likelihood difference scaling (Maloney & Yang, 2003; Knoblauch & Maloney, 2012). As in multidimensional scaling, our model assumes that each stimulus is represented in a subjective perceptual space where, in our case, the dimensions are color (C) and material (M). Rather than using a fixed location to represent each stimulus, we incorporate the idea that perception is noisy (e.g., Blakemore, 1990; Green & Swets, 1966) and model the representation of each stimulus as a bivariate Gaussian distribution. The mean of each Gaussian locates the corresponding stimulus in the perceptual space, while the covariance specifies the precision of the representation.

The model assumes that on each trial of the experiment, the actual representation of each stimulus (target and two tests) is a draw from the corresponding distribution and that the observer chooses the test stimulus whose representation on that trial is closest to that of the target. The probability that one test is chosen over another depends on the mean positions of their underlying representations, the magnitude of the perceptual noise, and a color-material weight This weight describes how differences along the two perceptual dimensions (color and material) are integrated when the observers select objects based on similarity.

In the model, we define the origin of the perceptual space by setting the position of the target to zero on each dimension. Similarly, we define the scale of the perceptual dimensions by setting the variance of the perceptual noise to one for each dimension. These conventions do not affect the model’s ability to account for the data. The observer’s performance is then described in terms of two key sets of parameters: (1) parameters that describe physical-to-perceptual mapping i.e., the mean position of each stimulus in the color-material perceptual space and (2) the color-material weight, which we denote as w. The computation of perceptual distances between the target and each test occurs only after distances on the color dimension have been scaled by *1-w* and distances along the material dimension have been scaled by 7-w. The color-material weight thus characterizes the relative importance of object color relative to object material in selection.

The model does include some substantive assumptions. First, we assume that positions along the color and material dimension are independent. That is, varying the position of a stimulus on the material dimension does not affect its position on the color dimension and varying the position of a stimulus on the color dimension does not affect its position on the material dimension. Second, we assume that the perceptual noise along the two underlying dimensions is independent. Third, we assume that the noise is additive and independent of the stimulus level. We return to consider the implications of these assumptions-in the Discussion.

We considered multiple variants of the perceptual model. These differed in two ways. First, we considered two different distance metrics (Euclidean and City-Block) for computing overall test-to-target distances, based on the weighted distances along each underlying dimension. Second, we considered four different ways of mapping nominal stimulus positions (labeled as -3 to +3 for each dimension, Figure 3) to the corresponding mean perceptual positions. In the Full variant, each non-zero nominal label was mapped onto its own mean perceptual position, so that 12 parameters were needed to describe to the mapping. In the Cubic, Quadratic, and Linear variants, the mean perceptual positions were obtained from cubic, quadratic and linear functions of the nominal labels (6, 4 and 2 parameters respectively) that pass through the origin (target coordinate of [0,0]). Thus 8 model variants were considered (2 metrics crossed with four positional-mapping variants). Details about the model implementation and how it was fit to the data are provided in Methods.

Our experimental design used Quest+, an adaptive trial selection procedure (Watson, 2017), together with the Euclidean/Cubic variant of our model. Given the parametric model, Quest+ selects for each trial the pair of test stimuli (7 levels per dimension, 49 possible stimuli, 1176 possible test stimulus pairs) that is predicted to yield the most information about the model parameters, given the selection data collected up to that point. The use of an adaptive method was critical for making the experiment feasible, as we estimate it would have taken ~40 hours per observer to measure the selection probabilities for all possible test pairs (~20 trials each for 1176 possible test pairs).

Development of the model and experimental procedures were guided by our findings in a preliminary experiment that used a subset of possible trial types. This experiment is described in a conference proceedings paper (Radonjić, Cottaris, & Brainard, 2018, also reviewed in Brainard, 2018 #13259).

### Experimental results

For each observer, we used a preregistered model selection procedure based on crossvalidated fit error to find which of the 8 model variants best accounted for each observer’s selection data. A detailed description of this procedure is available in Methods. We then used the best-fitting model variant to infer the positions of the stimuli in the perceptual color-material space and the color-material weight for each observer.

**Figure 4.**
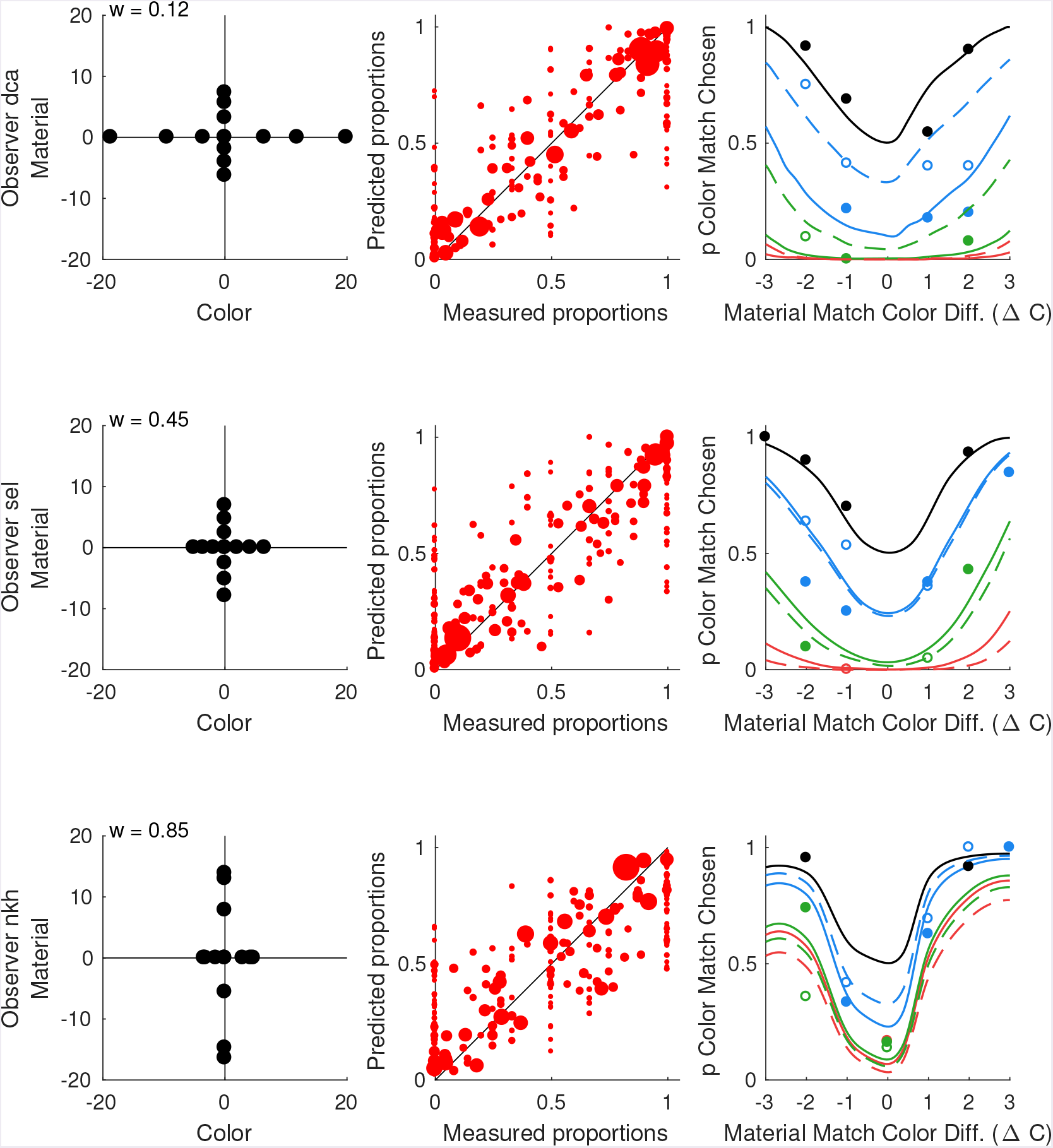
Example model solutions. Results are shown for the best-fitting model for three observers (top row: observer dca; middle row: observer sel; bottom row: observer nkh). **Left column**. Recovered mean perceptual positions for our test stimuli. The coordinates of the target stimulus are fixed at the origin. Non-target color levels are shown on the x-axis. They are ordered from C.3 on the left to C_+3_ on the right. Non-target material levels are shown on the y-axis. They are ordered from M_3 on the bottom to M_+3_ on the top. The inferred color-material weight isindicated at the top left. **Center column**. Measured selection proportions are plotted against proportions predicted by the model. The area of each data point is proportional to the number of trials run for the corresponding stimulus pair. One probability is plotted for each stimulus pair. Only data for stimulus pairs that were presented more than once are shown. **Right column**. Color-material trade-off functions predicted by the model solution (see main text). Measured probabilities for a subset of trials shown in the experiment are plotted against predicted probabilities. Symbol color matches the corresponding color-material trade-off function. Data corresponding to a prediction are plotted if more than 10 trials were run for the stimulus pair. Open circles correspond to dashed-lines (M_-3_, M_-2_, M_-1_) and filled circles correspond to solid-lines (M_+3_, M_+2_, M_+1_).

Figure 4 shows the model solution for three of our observers. These observers differ in their color-material weight (*dca* w = 0.12; *sel*, w = 0.46; *nkh*, w = 0.85). Each row shows data for one observer. The model solution is represented across three panels, which illustrate the recovered parameters, the quality of model fit, and what we refer to as color-material trade-off functions. Table 1 indicates which model variant was best for each observer.

**Table 1.** Results of model comparison. For each observer, the table shows the model variant that provided the best fit to the data. The same information is provided for the best positional variant of the model based on the alternative (non-best fitting) distance metric. The rightmost column indicates whether the cross-validation log-likelihoods obtained for the best fitting models for the two metric variants differed significantly. A paired-sample t-test, with α criterion level adjusted for multiple comparisons (one test for each observer: *p* = 0.05/12 = 0.0042) indicated that the two models differed significantly only for one observer (*cjz*, marked with asterisk). For four other observers p-values were smaller than 0.05 (as shown), but were not significant after the correction for multiple comparisons.

| Observer | Overall best model |  |  | Best model (alternative distance) |  |  |
| --- | --- | --- | --- | --- | --- | --- |
| initials | positions | distance | weight | positions | distance | weight |
| hmn | Full | Euclidean | 0.11 | Full | City-block | 0.22 |
| dca | Full | City-block | 0.12 | Quadratic | Euclidean | 0.14 |
| gfn | Cubic | Euclidean | 0.16 | Quadratic | City-block | 0.35 (p = 0.013) |
| lma | Quadratic | City-block | 0.37 | Quadratic | Euclidean | 0.43 |
| ofv | Full | City-block | 0.41 | Full | Euclidean | 0.46 |
| sel | Quadratic | City-block | 0.45 | Quadratic | Euclidean | 0.46 (p = 0.01) |
| ckf | Cubic | Euclidean | 0.51 | Quadratic | City-block | 0.64 |
| lza | Quadratic | City-block | 0.52 | Cubic | Euclidean | 0.82 (p = 0.03) |
| cjz | Cubic | City-block | 0.55 | Cubic | Euclidean | 0.86 (p < 0.001*) |
| jcd | Linear | City-block | 0.59 | Linear | Euclidean | 0.58 |
| nkh | Full | Euclidean | 0.85 | Full | City-block | 0.57 |
| nzf | Full | City-block | 0.88 | Full | Euclidean | 0.48 (p = 0.02) |

The left column of Figure 4 shows the inferred stimulus positions. The target is located at the origin. The x-axis shows the color dimension: points to the left of the origin indicate stimuli that are greener than the target and points to the right indicate stimuli that are bluer. The y-axis shows the material dimension: points below the origin indicate stimuli that are glossier than the target, while points above indicate stimuli that are more matte. Within each dimension, the mapping between nominal stimulus positions and perceptual positions is ordered (from C_-3_ on the left to C_+3_ on the right and from M_-3_ on the bottom to M_+3_ on the top). The inferred stimulus positions differ across observers. In addition, the inferred stimulus spacing along each dimension is not uniform. This should not be surprising, as without extensive preliminary experimentation there is no reliable way to choose stimuli that have uniform perceptual spacing for each observer.

The center column illustrates the quality of the model fit to the data. For each stimulus pair shown more than once, the measured proportion of trials one test was chosen relative to another is plotted against the corresponding proportion predicted by the model. The diagonal represents the identity line: the closer the points are to the diagonal, the better the agreement between model and data. The area of each plotted point is proportional to the number of trials run for a given stimulus pair: the larger the data point the more trials were shown. The model provides a reasonable account of the data, with the large plotted points lying near the diagonal.

The right column shows color-material trade-off functions. These are the model predictions for trials in which one of the tests is a *color match* and the other test is a *material match*. We use the term color match to refer to tests that have the same color as the target but differ in material, and the term material match to refer to tests that have the same material as the target but differ in color. The color-material trade-off functions show the proportion of time a color match is chosen (y-axis), when paired with the material matches. The color difference of the material match from the target is indicated on the x-axis. The black line shows the trade-off for a color match that is identical to the target (zero material difference: M_0_). When paired with the material match that is also identical to the target (zero color difference: C_0_), predicted selection proportion is at chance. As the color difference of the material match increases, the predicted probability that the observer chooses the color match increases and approaches 1. The red lines show the trade-off for a color match for which the material difference from the target is large (dashed line: M_-3_; solid line: M_+3_). When paired with the material match that is identical to the target (C_0_), the observer is predicted to select the material match (color match selection proportion near 0). As the color difference of the material match increases, however, the observer switches to selecting the color match, tolerating the difference in material. The green and blue lines indicate trade-off functions for intermediate values of color match material difference (small difference step in blue: M_-1_ is dashed and M_+1_ is solid line; medium difference step in green: M_-2_ is dashed and M_+2_, is solid line). These fall between the black and red lines. The relative steepness of the color-material trade-off functions reflects how readily the observer transitions to preferring the color matches over material matches. The steepness of the functions also varies across the three observers, and is qualitatively consistent with the differences in the inferred color-material weight. For example, the trade-off functions for the observer *nkh*, who has a high color-material weight (tends to make selection based on color), indicate very low tolerance for color differences of the material matches before the predicted selections switch to the color matches. The trade-off functions for observer *dca*, who has a low color-material weight are flattened, in comparison, indicating a large degree of tolerance for color differences of the material match. Note, however, that the trade-off functions depend both on the perceived spacing between the stimuli along the color and material dimensions, as well as on the color-material weight.

We evaluated the quality of the fit of the color-material trade-off functions to the data by plotting the measured selection proportions for the trials in which a color match test is paired with the material match. Because the trial selection was determined by the Quest+ procedure, only a subset of such trials was presented and the number of trials per pair varied across observers. Points are plotted only for pairs shown at least 10 times. Comparison of plotted points with corresponding prediction lines shows good agreement in most cases.

### Individual differences in color-material tradeoff

For each observer, the color-material weight inferred from the best-fitting model variant is shown in Figure 5 (green circles). There are large individual differences in the weights, which vary between 0.11 and 0.88. While half of the observers weight color and material roughly equally (weights between 0.41 and 0.59), for four observers selections are primarily guided by material (weights between 0.11and 0.37) while for the remaining two observers selections are primarily guided by color (weights greater than 0.8).

**Figure 5.**
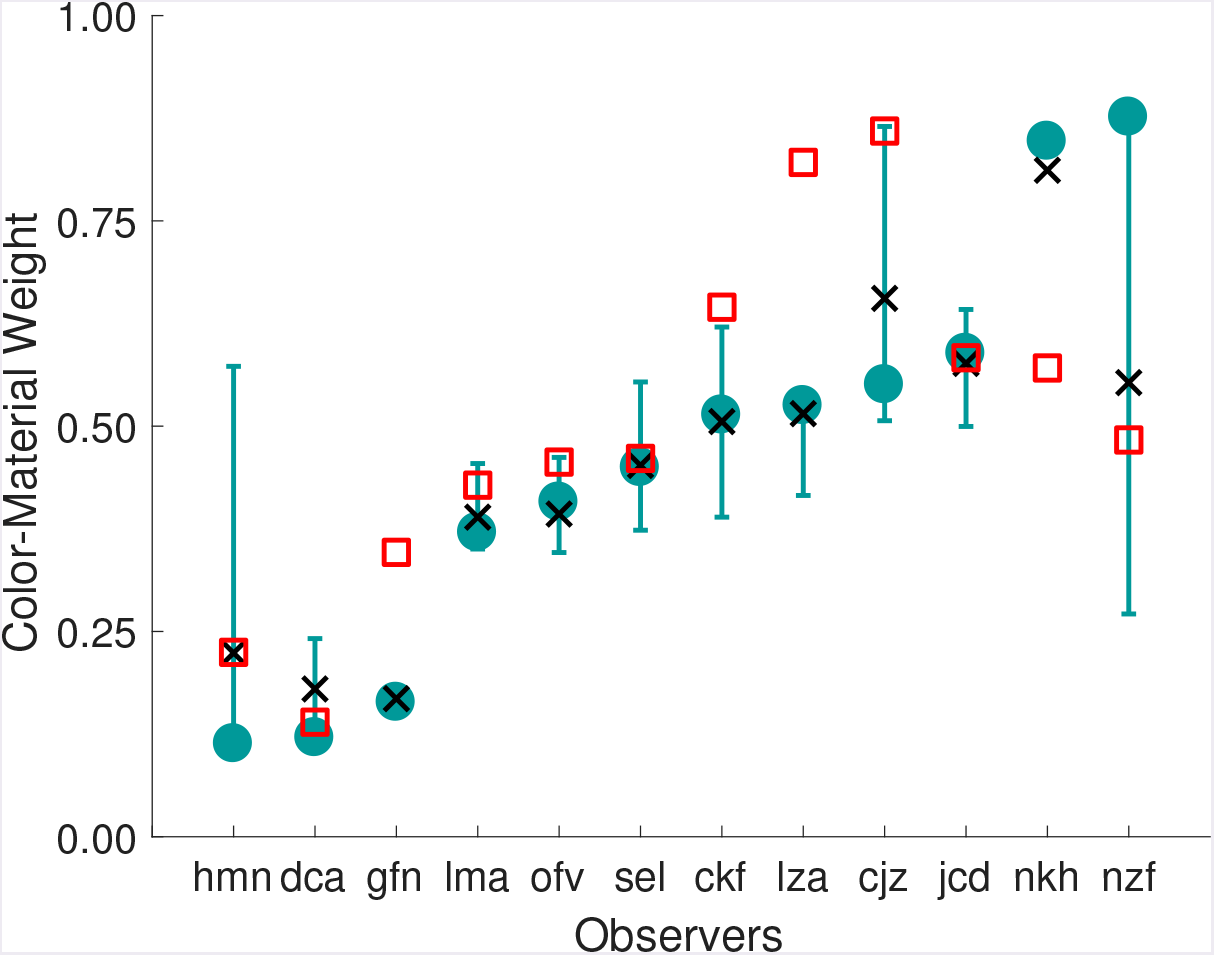
Color-material weight varies across observers. For each observer, the figure shows the color-material weight inferred from the best-fitting model variant (green circles). Which variant was best for each observer is provided in Table 1. Error-bars indicate the central 68% confidence interval (confidence range corresponds to the size of 1 SEM for a Gaussian distribution), obtained via bootstrapping. Black x symbols show the mean of the bootstrapped weights. Red squares show the weight inferred from the best-fitting model for the alternative distance metric.

We estimated the reliability of the color-material weight using a bootstrapping procedure (Efron & Tibshirani, 1993). For each observer, we resampled (with replacement) a new set of trials from the data and re-estimated the model parameters, using the best-fitting model for that observer. The error bars in Figure 5 show the central 68% confidence intervals from 100 bootstrap iterations. The black x symbols indicate the mean of the bootstrapped weights. For some observers the confidence intervals are small and the color-material weight is well-determined. For other observers the confidence intervals are large. We discuss this feature of the data in more detail below. Here, however, we note that the overall range of color-material weights remains large even if we consider only observers whose color-material weights have tight confidence intervals.

Table 1 provides a more detailed summary of the model fits. For each observer, the table indicates which positional (Full, Cubic, Quadratic or Linear) and metric (Euclidean or City-Block) model variant provided the best fit to the data. We also show which positional variant was best for the alternate (non-best fitting) distance metric. For 8 out of 12 observers the City-block metric provided the best fit, but the difference between City-block and Euclidean metrics was statistically significant for only 1 of 12 observers after correction for multiple comparisons (Table 1). Thus, our data do not strongly distinguish between the two distance metrics.

Across observers, the more complex positional variants tended to provide the best fits: there was only one observer for whom the Linear variant was selected as best. With the best-fitting distance metric, 5 observers’ data were fit best with the Full positional variant, 3 with the Cubic variant, and 3 with the Quadratic variant.

We also compared the color-material weights recovered using the two distance metrics. Figure 5 plots as red open squares the weights from the best-fitting positional model variant based the alternative distance metric. Across observers, the two weights are well correlated (Spearman rank-order correlation: *r*(10) = 0.78, *p* < 0.005), although for some observers the difference exceeds the confidence intervals on the weight inferred from the model based on the best-fitting metric. Moreover, a wide range of weights, indicative of large individual differences, is also observed for the best model based on the alternative metric.

### Limits to parameter identfiability

One of the key goals of our model was to independently describe the underlying stimulus representation in a perceptual color-material space and the color-material trade off. In other words, we aimed to uniquely determine (1) the parameters describing the stimulus positions and (2) the color-material weight. Two aspects of the results indicate that there are limits on how well this can be accomplished. First, some of the bootstrapped confidence intervals for the color-material weight are large (Figure 5). Second, examination of Figure 4 indicates the possibility of a systematic relationship between stimulus positions and the color-material weight. For the observer (*dca*) for whom the color-material weight is small, the inferred stimulus positions on the color dimension are expanded relative to those on the material dimension. For the observer (*nkh*) for whom the color-material weight is large, the opposite relation is seen: positions on the material dimension are expanded relative to those on the color dimension.

To investigate this further, we summarized the relation between the positions on the color and material dimension by first finding for each dimension the slope of the function mapping nominal stimulus position labels (integers between -3 and +3) to perceptual positions. We then computed the ratio of the slope for color to the slope for material. This *color-material slope ratio* is large when the positions on the material dimension are compressed relative to the positions on the color dimension (e.g., *dca*) and small when material is expanded relative to color (e.g, *nkh*). Thus, the color-material slope ratio provides an index for relative positional expansion on the two perceptual dimensions, and we can examine how it varies with the color-material weight.

Figure 6 shows the set of bootstrapped color-material slope ratios against the corresponding color-material weights, with results for each observer shown in a different color. The black open circles show the slope ratio and weight inferred from the best-fitting model applied to the complete data set for each observer. The figure illustrates that there is a systematic trade-off between the two aspects of the model solution. Within each observer, the higher the color-material weight, the lower the color-material slope ratio. The correlation between the two numbers is high statistically significant for every observer (Pearson correlation coefficients ranged from -0.99 to -0.91, *p* < 0.0001 for all observers). The distribution shown for each observer reflects the measurement uncertainty in determination of the two aspects of the solution. The confidence intervals shown in Figure 5 represent the central 68% of the x-axis variation for each observer shown in Figure 6.

**Figure 6.**
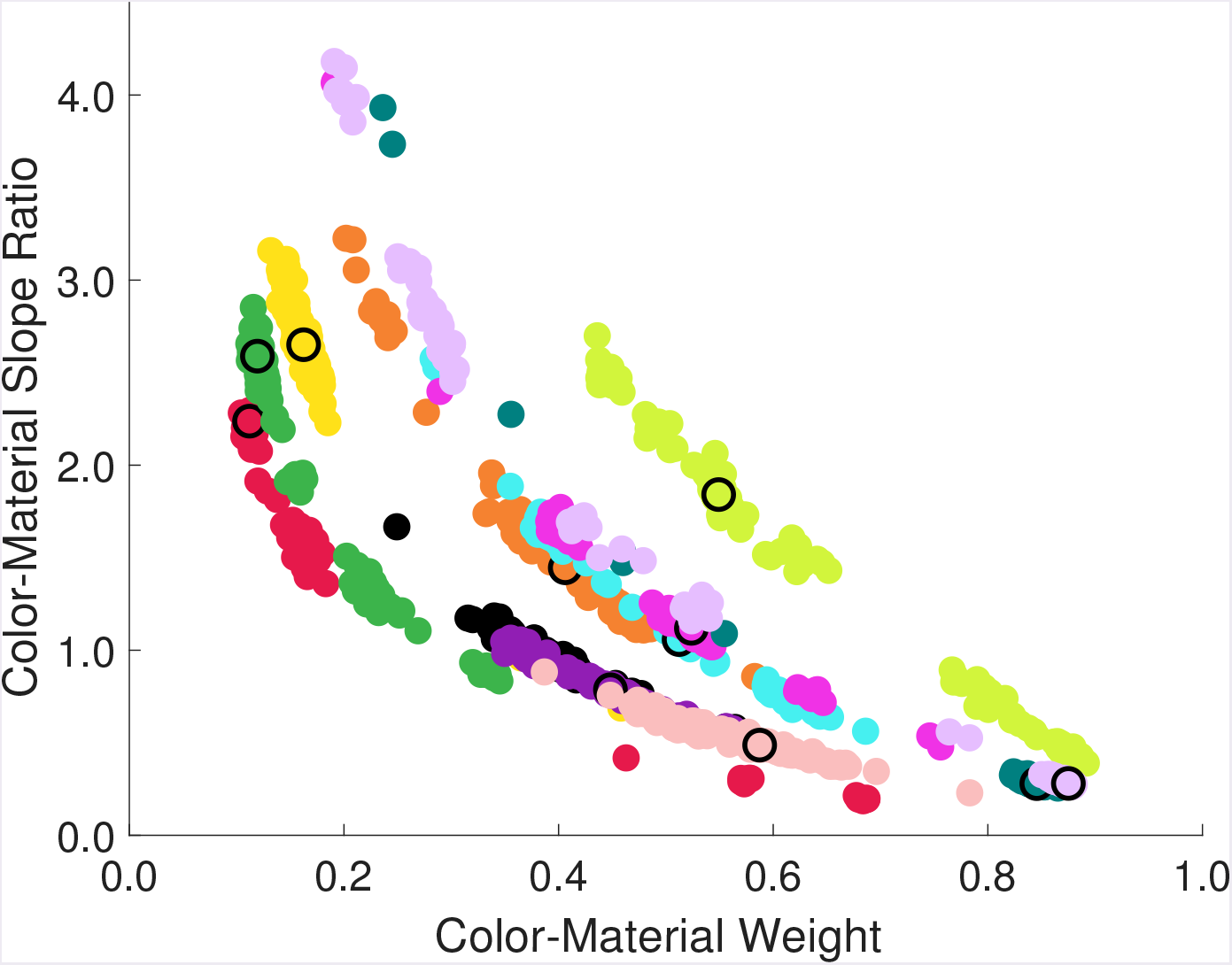
Color-material slope ratio is inversely related to the color-material weight. For each observer we plot the color-material slope ratio against the inferred weight for each bootstrap iteration (see text). Different observers are plotted in different colors. Black open circles plot the color/material slope ratio against the color-material weight inferred from the best-fitting model (full data set) and are superimposed over the corresponding observer’s data for different iterations.

Figure 6 demonstrates that stimulus positions and the color-material weight are entangled in the model solution: changes in color-material weight can be compensated by adjusting the stimulus positions without a large effect on the quality of the model fit. However, this trade-off between parameters is only partial: Figure 6 also illustrates clear individual differences across observers. First, the distributions of the bootstrapped color-material weight overlap minimally for some observers (e.g., green and yellow points vs. pink and lime points). Second, even when the range of color-material weights overlaps, the data for different observers can fall along distinct lines (e.g. red versus yellow points; pink versus lime points). This result emphasizes that there are clear individual differences in performance even for observers whose color-material weights are not differentiated by the data.

## Discussion

Color, material, texture and shape inform us about objects and guide our interactions with them. How vision extracts information about individual object properties has been extensively studied. Little is known, however, about how percepts of different properties combine to form a multidimensional object representation. Here we describe a paradigm we developed to study the joint perception of two different object properties, color and material. Our work builds on the literature on cue-combination, which also considers the multidimensional nature of object perception (Landy, Maloney, Johnston, & Young, 1995; Ernst & Banks, 2002; Hillis, Watt, Landy, & Banks, 2004; Knill & Saunders, 2003; Saarela & Olkkonen, 2017). What distinguishes our approach is that we move beyond threshold measurements to study supra-threshold differences (see also Ho, Landy, & Maloney, 2008; Hansmann-Roth & Mamassian, 2017).

On each trial of our object selection task observers viewed objects that vary in color and material (glossiness) and made selections based on overall similarity. We interpret the selection data using a novel observer model. The model allows us to describe the data in terms of the underlying perceptual stimulus representation and a color-material weight, which quantifies the trade off between object color and object material in selection. We find large individual differences in color-material weight across twelve observers: some observers rely predominantly on color when they select objects, others rely predominantly on material, and yet others weight color and material approximately equally.

Development of our observer model required us to overcome two fundamental challenges. The first arises because both the underlying perceptual representation of the stimuli and the way information is combined across perceptual dimensions are unknown and thus need to be recovered simultaneously. Although these two factors are conceptually different, their variation can have a qualitatively similar influence on the observers’ selection behavior. An important advance of our model is that it allows us to separate the contribution of the two factors. This separation works sufficiently well to allow us to establish that individual observers employ different color-material weights. At the same time, our work reveals limits on how precisely the contribution of the two factors can be separated. Improving the precision this separation represents an important direction for future work. We return to this point later in the Discussion.

The second challenge arises because as the number of dimensions studied increases and the stimulus range extends to include supra-threshold differences, the set of stimuli that could be presented grows far too rapidly for exhaustive measurement. This highlights the need for a theoretically-driven stimulus selection method, which would enable estimation of model parameters from a feasible number of psychophysical trials. To address this challenge, we implemented an adaptive stimulus selection procedure, which incorporated a seven-parameter variant of our model. The procedure is based on the Quest+ method (Watson, 2017) and selects on each trial the test stimulus pair that is most informative about the underlying model parameters. The strength of this approach is that it allows us to exploit appropriately complex models of how observers perform our task. One side-effect of using this efficient procedure, however, is that the power of the data to test the how well the model accounts for performance is reduced. We handled this by conducting a preliminary experiment as a part of model development (Radonjić, Cottaris, & Brainard, 2018). In this experiment we studied only a subset of stimulus pairs (color matches paired with material matches), using the method of constant stimuli, and showed that our model (Full positional variant with Euclidean distance metric) accounts well for the selection data.

As we noted above, our analyses show that in the model solution the recovered stimulus positions and color-material weight are not entirely independent: variation in the color-material weight can be compensated by variation in stimulus positions with minimal effect on the quality of the model’s fit. This trade-off in the recovered model parameters emerges because our model explicitly includes perceptual noise along each perceptual dimension. Counter-intuitively, this means that even when stimuli vary only along a single perceptual dimension (e.g. color), changing the stimulus spacing along that dimension need not have a large effect on observers’ predicted selection probabilities. Instead, such change in spacing can be compensated by changing the relative weight, which in turn affects then contribution of noise in the other (non-varied) dimension on the predicted selection probabilities. Although this compensatory effect is not complete, developing ways to more forcefully identify model parameters remains an important future research direction, as we and others move towards understanding the multidimensional nature of object representations. Below we outline three possible approaches to address this challenge.

The first approach is to obtain direct measurements of the underlying stimulus representations along each perceptual dimension. This could be achieved by conducting separate experiments in which observers are explicitly instructed to select objects based on only one aspect of the stimulus at a time (either color or material). This approach was taken in recent work that focuses on the independence of the perceptual representations of multiple object properties (Ho, Landy, & Maloney, 2008; Hansmann-Roth & Mamassian, 2017; Chadwick, Heywood, Smithson, & Kentridge, 2018; Rogers, Knoblauch, & Franklin, 2016; Qi, Chantler, Siebert, & Dong, 2015). It relies on the assumption that when instructed to attend to just one stimulus dimension, observers are able to ignore variations along the other dimensions. If the assumption holds, this approach combined with ours would provide additional power to recover stimulus positions along the attended dimension because it enables fitting the data without introducing the effects of noise along the non-attended dimension. There is no guarantee, however, that observers can or do strictly follow experimental instructions that direct them to attend only to a single perceptual dimension.

The second approach is to simplify our observer model and assume that the underlying stimulus representation is common across observers, so that any variation in performance is entirely due to the variation in the color-material weight. While the assumption that all observers perceive the stimuli in the same way may be questioned, it has a long history in the study of perception. Indeed, this approach is implicit in (1) efforts to develop standardized perceptual distance metrics and stimulus order systems (e.g., Brainard, 2003), (2) studies in which the conclusions about perceptual representations are based on data averaged across multiple observers (e.g., Shepard & Cooper, 1992, see Figure 2) and (3) work that employs multi-dimensional scaling to recover the common perceptual representation across observers together with the parameters that describe different weighting of the underlying dimensions by each observer (e.g., Carroll & Chang, 1970).

Finally, the third approach is to employ a multiplicative rather than an additive noise model. The precision of perceptual representations is often the highest at the current adaptation point (e.g., Loomis & Berger, 1979). This observation can be incorporated in the model by assuming that along each perceptual dimension noise scales as a function of test distance from the target. Reducing the noise near the perceptual axes might reduce the model’s ability to trade-off stimulus positions parameters and the weight. Along similar lines, one could consider modeling trial-by-trial variability as noisiness of the perceptual differences between stimuli, after the information has been combined across dimensions. Adding noise to the decision variable has been used in related work (Ramsay, 1977; Takane, 1978 #13208, see also Ho, Landy, & Maloney, 2008). Switching to a different noise model, however, would require careful consideration of whether the alternative model provides a better account of the data.

An additional assumption is that color and material are independent perceptual dimensions, so that variation in stimulus color has no effect on perceived glossiness and variation in stimulus glossiness has no effect on perceived color. The model embodies this assumption because the stimulus position along a perceptual dimension is a function of its nominal stimulus position along that dimension alone and is unaffected by the nominal stimulus position along the other dimension. For example, the stimuli that vary only in material relative to the target all have a color coordinate of 0 and vice versa. While this simplification provided a good starting point, it may be too restrictive. Ho et al. (2008) have shown that perceived glossiness varies with changes in the physical roughness of an object surface, and that perceived roughness varies with changes in physical glossiness, while Hansmann-Roth and Mamassian (2017) found a similar relationship between glossiness and perceived lightness. These findings suggest that such interactions might be present in our experiment (see also Qi, Chantler, Siebert, & Dong, 2015; Chadwick, Heywood, Smithson, & Kentridge, 2018; Rogers, Knoblauch, & Franklin, 2016).

The results we present here are based on experimental work that used a static object with single shape, one type of material change, and a limited range of variation in both color and material. Future research should expand the set of dimensions studied and the stimulus range along each dimension. In addition, the use of dynamic stimuli where observers compare objects as they rotate, each in a different phase, would minimize observers’ ability to make selections based on local image information (e.g., reflections in the specular highlights) and enforce similarity judgments based on global object appearance.

A key issue in the perception of object properties is how the visual system stabilizes its perceptual representations against variation in the conditions under which objects are seen, particularly against changes in the spectral and geometric structure of the illumination (Brainard & Radonjić, 2014; Fleming, 2017). It is of considerable interest to understand whether the way vision combines information about different classes of object properties is sensitive to viewing conditions. For example, perceived object color might be weighted less when objects are compared across changes in illumination spectrum. Similarly, perceived material might be weighted less when the comparisons are made across changes in the geometric properties of the illumination. By handling perceptual representations along multiple perceptual dimensions and by quantifying the relative importance of each dimension in performance, our methods and model extend naturally to the study of these questions.

## Methods

### Preregistration

Experimental design and the primary data analysis procedures for this study were preregistered before that start of the experiment. They are publicly available at: https://osf.io/263ua. Deviations from and additions to the preregistered plan are described in Appendix B.

### Apparatus

The stimuli were presented on a calibrated LCD color monitor (27-in. NEC MultiSync PA241W; NEC Display Solutions, Itasca, IL), in otherwise dark room. The monitors were driven at a pixel resolution of 1920 x 1080, a refresh rate of 60 Hz, and with 8-bit resolution for each RGB channel (NVIDIA GeForce GTX 780M video card; NVIDIA, Santa Clara, CA). The host computer was an Apple Macintosh with an Intel Core i7 processor. The experimental programs were written in MATLAB (MathWorks; Natick, MA) and relied on routines from the Psychophysics Toolbox (Brainard, 1997; Pelli, 1997, http://psychtoolbox.org) and mgl (http://justingardner.net/doku.php/mgl/overview).

In the experiment, the observer’s head position was stabilized using chin cup and forehead rest (Headspot, UHCOTech, Houstion, TX). The observer’s eyes were centered horizontally with respect to the display, and centered vertically approximately 2/3 of the distance from the bottom to the top of the display. The observers viewed the stimuli binocularly. The distance from observer’s eyes to the monitor was 75 cm.

### Task

On each trial of the experiment, three images were displayed on the monitor (Figure 2). Each image was a rendering of a stimulus scene consisting of a room with a blob-shaped object in the center. The object in the middle scene was the target object, while the objects in the left and in the right scenes were the test objects for the trial. Observers were instructed to “select the test object that is most similar to the target”. They indicated their choice by pressing a button on a game controller and were allowed to take as much time as they needed to respond. After they responded, a small black dot briefly flashed above the test object that was selected, indicating to the observer that his/her response has been recorded. The next trial started after a brief, variable interstimulus interval (ranging from 0.63 s to 1.51 s with the mean of 1.04 s; see experimental procedures).

### Stimulus scenes

We rendered 49 stimulus scenes. These were identical except for the surface reflectance of the blob-shaped object. Each scene was modeled as a room, whose walls were gray (assigned the diffuse surface reflectance of the Macbeth color checker chart sample from row 4, column 2; http://www.babelcolor.com/colorchecker.htm). The floor consisted of white and dark-gray tiles arranged in the checkerboard pattern (Macbeth chart sample reflectances: row 4, columns 1 and 4, respectively). A gray square pedestal was positioned in the middle of the room (Macbeth chart sample reflectance: row 4, column 2), with the blob-shaped object placed on top of it.

The blob-shaped object was generated from an icosahedral mesh that approximated a sphere. Each side of the mesh was subdivided into 64 facets and a sinusoidal perturbation was added to the x, y and z coordinates of each facet vertex. A different sinusoid was used for each (x, y, z) dimension. Surfaces in the room were modeled as matte (diffuse surface scattering model in the Mitsuba renderer). The one exception was a blob-shaped object for which we used Mitsuba’s anisotropic Ward model. Further details on the rendering procedures are provided below.

The room was illuminated by an area light, which covered the entire surface of the white ceiling (Macbeth chart reflectance sample: row 4, column 1). The illumination spectrum was set to a CIE daylight with 6700 K correlated-color temperature (D67). The illumination was fixed across all stimulus scenes.

### Blob-shaped object manipulation

Across stimulus scenes we varied two surface properties of the blob-shaped test objects: their color (i.e., diffuse surface reflectance) and their material, specifically glossiness. The test objects varied along 7 different levels on each dimension yielding total of 49 different stimulus scenes (i.e., 49 different test objects). The target object (center image in Figure 1) was positioned in the center of this stimulus space: its color was greenish-blue (color coordinate denoted as C0) and its material was semi-glossy (material coordinate denoted as M0). The test objects could vary relative to the target in color only, material only or both. We also included a test object that was identical to the target.

The variation in the color dimension was achieved by varying test object’s diffuse surface reflectance across 7 different levels. This resulted in a variation in test color appearance from greenish to bluish. Under the scene illumination (D67) the xy coordinates of test objects were: [0.247, 0.311] for C_-3_ (most green-appearing), [0.244, 0.305] for C_-2_, [0.242, 0.298] for C._1_, [0.239, 0.292] for C_0_ (equal to the target color level), [0.238, 0.287] for C_+1_, [0.236, 0.281] for C_+2_ and [0.235, 0.276] for C_+3_ (most blue-appearing). These values are computed assuming flat matte surfaces of corresponding surface reflectance functions under diffuse illumination. Across levels, the relative luminance of the test object under fixed scene illumination decreased gradually as the reflectance changed (so that the luminance of the greenest test C.3 was approximately 11% higher than that of the bluest test C_+3_; see right column in Figure 2).

The spectral reflectance was set to 0.28 for all test objects (details on the rendering algorithm are available below). The variation in material (glossiness) of the test object was achieved by varying surface parameters α_U_ and α_V_ (in Mitsuba renderer notation). These parameters control anisotropic roughness of the material along the tangent and bitangent directions of reflected light (Wenzel, 2014), i.e., the amount of blurring at the specular lobe (Ho, Landy, & Maloney, 2008, p. 197): low α_U_ and α_V_ values correspond to smooth, glossy-appearing surfaces, while high values correspond to matte-appearing surface with rough microstructure. For each test, α_U_ and α_V_ were set to the same value. For the target glossiness level (M_0_) α_U_ and α_V_ were set to 0.1. The values for the remaining test were: 0.035 for M_-3_, 0.055 for M_-2_, 0.08 for M_-1_, 0.14 for M_+1_, 0.17 for M_+2_ and 0.02 for M_+3_.

We adjusted the levels of variation in color and material based on observations we made in the preliminary experiment (Radonjić, Cottaris, & Brainard, 2018). Our aim was to select the stimulus spacing that would produce gradual shifts in performance as the tests varied along each dimension. In the color dimension, the spacing between adjacent levels was fixed to 2.61 △E units in the CIELAB color space, which is approximately uniform. This spacing is slightly smaller 1 just noticeable difference (which is estimated to be 3.6 CIELAB ΔE Brainard, 2003, p.203). CIELAB coordinates were computed from the target reflectance and scene illumination spectra, assuming a matte flat surface under uniform illumination and using the scene illumination’s XYZ value as the white point for conversion. To determine the surface reflectances of the remaining (non-target) color levels that correspond to the desired spacing (anc test CIELAB/XYZ values) we used standard colorimetric methods (Brainard & Stockman, 2010) and a three-dimensional linear model for surface reflectance functions derived from analysis of the spectra of Munsell papers (Nickerson, 1957, http://psychtoolbox.org, first three columns of matrix B_nickerson).

The data we collected in the preliminary experiment suggested that the spacing in the material dimension was larger than optimal. We therefore reduced the material step size by eliminating the most glossy (α_U,V_ = 0.007) and the most matte (α_U,V_ = 0.4) levels (from those used in the preliminary experiment). We further fine-tuned the spacing on the material dimensio so that the within-dimension discrimination between adjacent material levels (for fixed, target color) was roughly the same level of difficulty as the discrimination between adjacent color levels. This adjustment was based on informal judgments by authors AR and DHB. Note, however, that our attention to spacing was only for the purpose of maximizing the power of the data. Our model (which we describe below in detail) does not assume that the spacing between our stimuli is uniform within dimension or equated across dimensions.

### Stimulus generation

The stimulus scenes were modeled in Blender, an open-source 3-D modeling and animation package (https://www.blender.org/) and rendered using Mitsuba, a physically-realistic open-source rendering system (Wenzel, 2014, https://www.mitsuba-renderer.org). For the rendering, we used a bidirectional path tracer integrator, which included modeling of indirect illumination (up to 6 bounces) and a low discrepancy sampler (number of samples 1024). Rendering was facilitated via RenderToolbox3 routines (Heasly, Cottaris, Lichtman, Xiao, & Brainard, 2014), which enabled us to assign and manipulate surface reflectance properties of objects in the scene. Code that implements the renderings is publically available (see below) and includes all rendering materials (surfaces, illuminations, renderer specification, etc).

Each rendered image was a 31-plane hyperspectral image, specified at 10 nm wavelength sampling between 400 and 700 nm. We converted each image into a three-plane LMS image by computing the pixel-by-pixel excitations that would be produced in the human L-, M-, and Scones, using the Stockman-Sharpe 2-degree cone fundamentals (CIE, 2007, Stockman & Sharpe, 2000). The LMS images were then converted into RGB images for presentation based on the monitor calibration and standard methods (Brainard, Pelli, & Robson, 2002). Monitor calibration measurements were made using a PR-650 SpectraScan radiometer (Photo Research Inc, Syracuse, NY). They included the spectral power distributions of the monitor RGB primaries as well as their gamma functions. Finally, all RGB rendered images were scaled by a common factor to maximize the fraction of the display gamut used by the stimulus set. This last image manipulation is equivalent to increasing the illumination irradiance by a fixed factor across all stimulus scenes.

### Proximal stimulus

At the 75 cm viewing distance, the size of each stimulus image was 13.7° x 10.3° of visual angle (18 x 13.5 cm). The size of the blob-shaped object was 6.2° x 5.95° of visual angle (8.14 x 7.8 cm). Mean stimulus image luminance, averaged across 49 stimulus images, was 43.12 cd/m^2^ (*SD* = 0.29; mean value for each image was computed by averaging across pixels). The luminance of the black background that images were presented against was 0.96 cd/m^2^.

### Observers

Twelve observers participated in the experiment (8 female, 4 male; age 20-55; mean age 28.7). All observers had normal or corrected-to-normal vision (20/40 or better in both eyes, assessed using Snellen chart) and normal color vision (0 Ishihara plates read incorrectly, Ishihara, 1977). One additional observer (male, age 34) was recruited but excluded from the experiment before completion due to non-compliance with experimental instructions (pressing response buttons randomly, leaving the laboratory in the middle of an experimental block) and unreliability (repeatedly missing scheduled experimental sessions). Data from this observer were not analyzed. This exclusion criterion was not explicitly described in our pre-registration.

We did not conduct any formal analysis to determine the number of observers or number of trials per observer. In our preliminary experiment (Radonjić, Cottaris, & Brainard, 2018), we found large individual differences in color-material weighting across 5 observers. We judged that a sample size of 12 observers would allow us to complete data collection within a manageable time period (6-8 weeks) while collecting enough data to obtain a reasonable estimate of the variation in color-material weights across individuals. Both the planned number of observers and the planned number of trials per observer were specified in the pre-registration. All experimental procedures were approved by University of Pennsylvania Institutional Review Board and were in accordance with the World Medical Association Declaration of Helsinki.

### Experimental design

Each experimental block consisted of 270 trials, divided into 9 sub-blocks of 30 trials each. In 8 out of these 9 sub-blocks, the stimulus pairs presented on each trial were determined using the adaptive Quest+ procedure (Watson, 2017, Matlab implementation available at: https://github.com/BrainardLab/mQUESTPlus). In the remaining sub-block, the test pairs were determined by sampling randomly from the stimulus set.

Implementing Quest+ procedure requires specifying the model underlying observers’ selections and the parameter space corresponding to this model. Across trials, Quest+ selects the stimulus pair for each trial that would be most informative for determining the underlying model parameters, given the observer’s selection data prior to that trial.

We implemented Quest+ over a 7-dimensional parameter space, which corresponds to the cubic version of our model (which we describe in detail below). One parameter is the color-material weight. Three parameters map the nominal positions of the stimuli along the physical color dimension to their positions along an underlying perceptual color dimension. Three additional parameters map the nominal positions of the stimuli along the physical material dimension to their positions along an underlying perceptual material dimension. For both color and material, the three parameters are linear, quadratic and cubic coefficients for a cubic polynominal.

In our implementation of Quest+, the parameters were allowed to vary over a range of linearly spaced values, which we determined based on the results of our preliminary experiment (Radonjić, Cottaris, & Brainard, 2018). For both color and material mappings these ranges were: from 0.5 to 6 (5 levels) for linear coefficient, from -0.3 to 0.3 (4 levels) for the quadratic coefficient, and from -0.3 to 0.3 (4 levels) for the cubic coefficient. For the color-material weight the range was from 0.05 to 0.95 (5 levels). As in our model implementation (described below) we also constrained the parameters so that the stimulus positions in the perceptual space vary monotonically within the range -20 to +20 and we enforced a minimal spacing of 0.25 between the adjacent test positions (which is equal to ¼ of representational noise). We used a lookup table to finding the likelihood of observer’s response given a test stimulus pair and any set of model parameters. This lookup table was based on the Euclidean distance metric and was the same as the one used for modeling the selection data.

In 6 out of 8 Quest+ sub-blocks, the stimulus pairs that could be shown on each trial were sampled adaptively over the full stimulus range (−3 to +3, i.e.: C_-3_ to C_+3_ and M_-3_ to M_+3_). To ensure some diversity in stimulus selection across trials, we also included 2 sub-blocks in which Quest+ was run over a restricted stimulus range (one sub-block used the -2 to +2 stimulus range, while the other used a -1 to +1 stimulus range). Trials were run in groups of 9, with one trial from each sub-block chosen in random order before moving on to the next group of 9 trials.

The Cubic model based on Euclidean distance that was used to drive Quest+ represents only one variant of our model suite. It was not possible to know *a priori* which model variant would best describe the data for a given observer. We implemented the cubic model in our trial-by-trial stimulus selection. This was the most complex model we could feasibly implement while keeping the time between trials (needed to compute the next most informative stimulus pairs) reasonably short (~ 1s). The choice of Euclidean (rather than City-block) distance metric was arbitrary.

### Experimental procedures

At the beginning of the first experimental session, observers listened to detailed experimental instructions and completed a brief set of training trials. The full text of the instructions, which describe the task and experimental procedures (including how to respond, when and how to take a break within a block of trials, etc) is available in Appendix A. The training set consisted of 4 experimental trials in which the target was paired with one test that was identical to it (C0M0) and another that differed from it either only in color or only in material by the largest difference step (stimuli: C_-3_M_0_, C_+3_M_0_, C_0_M_-3_, C_0_M_+3_). The training data was not analyzed.

Each observer completed 8 experimental blocks of trials (270 trials per block). Typically, observers completed the blocks in 4 experimental sessions, each done on a different day. Within a session, which was approximately one-hour long, observers completed two blocks of trials, separated by a 5-10 minute break. An exception was observer *hmn* (male, age 32) who completed the experiment in 8 sessions (1 experimental block per hour-long session).

### Code

Model code is publicly available at https://github.com/BrainardLab/BrainardLabToolbox, (ColorMaterialModel directory). Experimental code is publicly available at https://github.com/BrainardLab/ColorMaterial.

## Model implementation and selection

### Computation of perceptual distances

We compute perceptual distances using either the Euclidean or City-block metric. With the Euclidean metric, the distance between the target *T* and the test *T_1_* (*d_T-T1_*) on each trial is computed as:

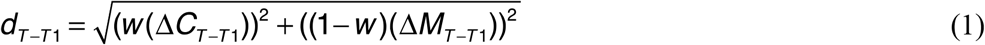

while the distance between the target *T* and the test *T_2_* (*d_T-T2_*) is computed as:

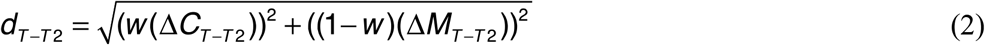

With the City-block metric version of the model, the corresponding formulas are:

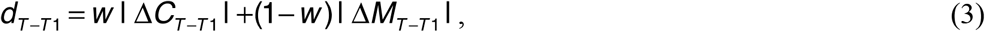

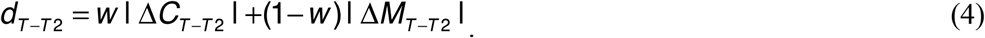

In the equations above, *w* denotes the color-material weight, ΔC denotes the distance between the target and a given test along the perceptual color dimension and △M denotes the distancebetween the target and a given test along the perceptual material dimension. On a given trial, the observer selects test T_1_ if *d_T-T1_ < d_T-T2_*, and T_2_ otherwise.

### Model origin and scale

Our model employs two conventions that define the units for the underlying perceptual dimensions. First, we define the target position as the origin of the perceptual space. Second, we model the zero-mean Gaussian perceptual noise for each stimulus as having a standard deviation of one along each perceptual dimension. This assumption defines the scale of the perceptual dimensions, with the mapping between physical and perceptual positions then fit with respect to this scale. In the model implementation, for each dimension we restricted the range over which stimulus positions can vary to -20 to +20 and we enforced the minimum spacing between adjacent stimuli to be at 0.25 (¼ of the standard deviation of the representational noise).

### Model variants

The full variant of our model (for either choice of distance metric) has 13 free parameters: the color-material weight *w*, 6 positions on the color dimensions that correspond to the 6 non-target color levels and 6 positions on the material dimension that corresponds to 6 nontarget material levels. We use numerical search to find the weight and the positions that best fit each observers’ selection data in a maximum likelihood sense.

Conducting the numerical search required us to be able to compute the likelihood of an observer’s responses for any pair of tests, given the color-material weight and the positions of the two tests in the underlying perceptual color-material space. We do not have an analytic formula for computing these likelihoods. We therefore pre-computed them using forward Monte Carlo simulation and stored them in a gridded multi-dimensional lookup table. More specifically, we constructed a 5-dimensional lookup table with dimensions: (1) color-material weight, (2) perceptual color coordinate of the first test, (3) perceptual material coordinate of the first test, (4) perceptual color coordinate of the second test, (5) perceptual material coordinate of the second test. For the color-material weight, we linearly sampled 10 grid weight values between 0 and 1. For color and material dimensions we linearly sampled 20 grid values between -20 and +20 corresponding to perceptual positions on each dimension. We simulated 3000 trials of test comparisons for each combination of parameters in the grid, and averaged over these to obtain the likelihood of each test being chosen when paired with each other test (for any sampled weight value). This table was used to estimate the likelihood of each response for any pair of stimulus positions within a predefined range (−20 to 20) and any weight value, using cubic interpolation (via Matlab’s griddedInterpolant function). We constructed two separate lookup tables, one based on the Euclidean distance metric and one based on the City-block distance metric.

For each metric, we compared four variants of our model, which differed in the complexity of the mapping between physical and perceptual stimulus positions. The full variant, described above, had 13 free parameters. In the simpler variants, we constrained the mapping between physical and perceptual stimulus positions along each dimensions to have a parametric form, with perceptual positions being described as a linear, quadratic, or cubic function of nominal stimulus positions (thus requiring 2, 4 or 6 free positional parameters, respectively, in addition to the color-material weight). Because the nominal position of target object color and material were 0 and mapped onto the [0,0] coordinate in the perceptual color-material space we did not include an affine term in our linear, quadratic or cubic mappings.

For each observer and choice of distance metric we used 8-fold cross-validation to compare the full model variant to three simpler (more constrained) variants. We evaluated the models by comparing the average cross-validated log-likelihoods of the fits across 8 crossvalidation partitions using a paired-sample t-test. For each partition, model parameters were determined from 7/8 of the trials from the full data set and the cross-validated log-likelihood was evaluated on the remaining 1/8 of the trials. We started the comparisons from the full model, which is the most complex and we asked if the cross validation log-likelihood was significantly higher for the full than for the cubic model (using the α-level of 0.05 for a one-tailed test). If it was, we concluded that full model best accounted for the data. Otherwise, we eliminated the full model from consideration and proceeded to compare log-likelihoods for the cubic and quadratic model, using the same method and criterion. If cubic model was significantly better than the quadratic, we concluded that cubic model best accounted for the data. Otherwise, we eliminated cubic model and continued to compare the quadratic and linear models using the same procedure. We followed this procedure separately for each observer to establish the model that best accounted for this observer’s selection data, given the choice of metric.

We conducted this procedure for both Euclidean and City-block metric separately. In the comparison for a given observer we used the same cross-validation data partition both across model variants and across different metrics. A different partition was used for each observer. To select best overall model, we compared the mean cross-validation log-likelihood of the best-fitting Euclidean-based and the best-fitting City-block-based model and selected the model that had the highest average cross-validated log-likelihood.

The model comparison procedures described above are conducted following the preregistered analysis plan. To examine whether the log-likelihoods of the two best-fitting models based on City-block-distance and Euclidean-distance differed significantly from each other, we also compared their average cross-validated log-likelihoods using a paired t-test (two-tailed, with α criterion level adjusted for multiple comparisons, one test for each observer: *p* = 0.05/12 = 0.0042) consider this to be a post-hoc analysis (as it was not described in the pre-registration document; see Appendix B), but never-the-less useful for understanding how model goodness-of-fit depends on the underlying metric. Thus we report the result of this comparison in addition to specifying which model had the lowest average cross-validated log-likelihood.

## Appendix A. Experimental instructions

Below we provide experimental instructions verbatim (text in italics). These were read to the observer at the beginning of the first experimental session. At the beginning of each following section the observers were only reminded of the task (experimenter only read the paragraph that begins with “your task”).

*“During the experiment we will ask you to view the stimuli from the position indicated by this chin rest. At this time, please adjust the chair so that your chin is positioned at the chin rest, your forehead is leaning against the upper bar and you are sitting comfortably with your back straight. We will describe your task in the experiment using a couple of training trials. “*

The experimenter starts the training, which consists of four trials (i.e., two identical illuminant-constant trials, repeated twice; competitors in the first and third trial are C_1_ and C_4_; in the second and and forth C_2_ and C_5_). Training data is not analyzed.

*“On each trial three scenes will be presented on the screen, each containing one object. τhe object in the middle scene is the target for the trial. τhe objects in the left and in the right scene are the test objects.”*

*“Your task is to select the test object that is most similar to the target. On some trials, both tests will be very similar to the target. Always try to pick the one that is most similar. “ “You will use this joystick to provide the response. To select the test on the left, press button 1 on the left. To select the test on the right, press button 3 on the right. When you make the choice a small black dot will briefly flash just above the test object you selected. At this point the trial will end and the new set of objects will appear on the screen.”*

*“There are 270 trials within a block and they are divided into 9 sets of trials. After each set, you will have a chance to take a break if you need one. The experiment will pause and the voice will inform you how many sets in this block are done and how many are left. You can take as much time as you need to rest between the sets. When you want to continue, press one of the front buttons on the joystick and the next trial will start. After a block of trials is finished, the screen will turn gray. At that point, we will take a 5-10 minute break before we continue with the second block of trials. Today we will do 2 blocks of trials.”*

*“It is very important that your head is positioned at the chin rest while you are viewing the stimuli. Please make sure you remain in this position during the set of trials. You will be able to move, stretch or adjust your position in between the sets or between the blocks. Do you have any questions?*

*“I will now initiate the experiment and leave the room. The experiment will start shortly after.”* Experimental program pauses for 15 seconds before the first trial to allow experimenter to leave the room.

## Appendix B. Deviations from and additions to preregistered plan for data collection and analysis

Below we summarize deviations from the preregistered plan for data collection and analysis, as well as list additional exploratory analyses reported here but that were not included in the preregistered plan.

Changes in experimental procedures:

- One observer was recruited but excluded from the experiment before completion due to noncompliance with experimental instructions (see Methods). Data from this observer were not analyzed.

Additional (exploratory) data analyses, not included in the preregistered data analysis plan:

- We plot color-material trade-off functions and data from the subset of trials in which the tests were a material match and a color match to describe the quality of the model fit to the data. Color-material trade-off functions and the corresponding data were used in this way for the preliminary experiment (Radonjić, Cottaris, & Brainard, 2018).
- We compared the quality of the fit of the two best models based on two distance metrics by conducting a 2-tailed paired t-test on cross-validated log-likelihoods. We also quantified the general agreement between the color-material weights inferred from the best models based on different distance metrics by computing a Spearman rank-order correlation coefficient.
- To investigate the relationship between the parameters describing the stimulus representation and the color-material weight, we used the bootstrapped data to compute color-material slope ratios and compare them to recovered color-material weight. We illustrate this relationship both graphically (Figure 6) and quantitatively (Pearson correlation coefficients).

## Footnotes

1 For expository purposes, we use the term *material* to refer to the physical glossiness, which is a function of the geometric reflectance properties of object surfaces. Similarly, we use the term *color* to refer to the diffuse surface spectral reflectance of objects. When it is not clear from context, we will explicitly distinguish the perceptual correlates of these physical properties (e.g., *perceived material* and *perceived color)*.

## References

Blakemore, C. (1990). Vision: Coding and Efficiency. Cambridge: Cambridge University Press.

Brainard, D. H. (1997). The Psychophysics Toolbox. Spatial Vision, 10(4), 433–436.

Brainard, D. H. (2003). Color appearance and color difference specification. In S. K. Shevell (Ed.), The Science of Color (2 ed., pp. 191–216). Oxford: Optical Society of America; Elsevier Ltd.

Brainard, D. H., Cottaris, N. P., & Radonjić, A. (2018). The perception of colour and material in naturalistic tasks. Interface Focus, 8(4) http://rsfs.royalsocietypublishing.org/content/8/4/20180012.abstract.

Brainard, D. H., Pelli, D. G., & Robson, T. (2002). Display characterization. In J. P. Hornak(Ed.), Encyclopedia of Imaging Science and Technology (pp. 172–188). New York, NY: Wiley.

Brainard, D. H., & Radonjić, A. (2014). Color constancy. In Werner J. S. & L. M. Chalupa (Eds.), The New Visual Neurosciences (pp. 545–556). Cambridge, MA: MIT Press.

Brainard, D. H., & Stockman, A. (2010). Colorimetry. In M. Bass, C. DeCusatis, J. Enoch, V. Lakshminarayanan, G. Li, C. Macdonald, V. Mahajan & E. van Stryland (Eds.), The Optical Society of America Handbook of Optics, 3rd edition, Volume III: Vision and Vision Optics (pp. 10.11–10.56). New York: McGraw Hill.

Carroll, J. D., & Chang, J.-J. (1970). Analysis of individual differences in multi dimensional scaling via an n-way generalization of “Eckart-Young” decomposition. Psychometrika, 35(3), 283–319. https://doi.org/10.1007/BF02310791.

Chadwick, A. C., Heywood, C. A., Smithson, H. E., & Kentridge, R. W. (2018). Beyond scattering and absorption: Perceptual un-mixing of translucent liquids. Journal of Vision, 18

Chadwick, A. C., & Kentridge, R. W. (2015). The perception of gloss: A review. Vision Research, 109, 221–235.

Efron, B., & Tibshirani, R. J. (1993). An introduction to the bootstrap. New York: Chapman and Hall/CRC

Ernst, M. O., & Banks, M. S. (2002). Humans integrate visual and haptic information in a statistically optimal fashion. Nature, 415, 429. http://dx.doi.org/10.1038/415429a.

Fleming, R. W. (2014). Visual perception of materials and their properties. Vision Research, 94, 62–75. http://www.sciencedirect.com/science/article/pii/S0042698913002782.

Fleming, R. W. (2017). Material Perception. Annual Review of Vision Science, 3(1), 365–388. https://doi.org/10.1146/annurev-vision-102016-061429.

Foster, D. H. (2011). Color constancy. Vision Research, 51, 674–700.

Green, D. M., & Swets, J. A. (1966). Signal detection theory and psychophysics. New York: John Wiley & Sons.

Hansmann-Roth, S., & Mamassian, P. (2017). A Glossy Simultaneous Contrast: Conjoint Measurements of Gloss and Lightness. i-Perception, 8(1), 2041669516687770. https://doi.org/10.1177/2041669516687770.

Heasly, B. S., Cottaris, N. P., Lichtman, D. P., Xiao, B., & Brainard, D. H. (2014).RenderToolbox3: MATLAB tools that facilitate physically based stimulus rendering for vision research. Journal of Vision, 14(2) http://www.journalofvision.org/content/14/2/6.abstractN2

Hillis, J. M., Watt, S. J., Landy, M. S., & Banks, M. S. (2004). Slant from texture and disparity cues: optimal cue combination. Journal of Vision, 4(12), 967–992.

Ho, Y. X., Landy, M. S., & Maloney, L. T. (2008). Conjoint measurement of gloss and surface texture. Psychol Sci, 19(2), 196–204.http://www.ncbi.nlm.nih.gov/entrez/query.fcgi?cmd=Retrieve&db=PubMed&dopt=Citation&listuids=18271869.

Ishihara, S. (1977). Tests for Colour-Blindness. Tokyo: Kanehara Shuppen Company, Ltd.

Knill, D. C., & Saunders, J. A. (2003). Do humans optimally integrate stereo and texture information for judgments of surface slant? Vision Research, 43(24), 2539–2558. http://www.sciencedirect.com/science/article/pii/S0042698903004589.

Knoblauch, K., & Maloney, L. T. (2012). Modeling Psychophysical Data in R (Use R!). New York: Springer.

Koenderink, J., & van Doorn, A. J. (2004). Shape and Shading. In L. Chalupa & J. S. Werner.(Eds.), The Visual Neurosciences. (Vol. 2, pp. 1090–1105). Cambridge, MA: MIT Press.

Kruskal, J. B. (1964). Multidimensional scaling by optimizing goodness of fit to a nonmetric hypothesis. Psychometrika, 29, 1–27.

Landy, M. S., Maloney, L. T., Johnston, E. B., & Young, M. (1995). Measurement and modeling of depth cue combination: in defense of weak fusion. Vision Research, 35(3), 389–412.

Loomis, J. M., & Berger, T. (1979). Effects of chromatic adaptation on color discrimination and color appearance. Vision Research, 19, 891–901.

Maloney, L. T., & Yang, J. N. (2003). Maximum likelihood difference scaling. Journal of Vision, 3(8), 573–585. http://www.journalofvision.org/content/3/8/5/.

Nickerson, D. (1957). Spectrophotometric data for a collection of Munsell sample.. Washington, DC: U.S. Department of Agriculture.

Olkkonen, M., & Ekroll, V. (2016). Color Constancy and Contextual Effects on Color Appearance. In J. Kremers, R. C. Baraas & N. J. Marshall (Eds.), Human Color Vision (pp. 159–188). Cham: Springer International Publishing.

Pelli, D. G. (1997). The VideoToolbox software for visual psychophysics: transforming numbers into movies. Spatial Vision, 10(4), 437–442.

Qi, L., Chantler, M. J., Siebert, J. P., & Dong, J. (2015). The joint effect of mesoscale and microscale roughness on perceived gloss. Vision Research, 115, 209–217. http://www.sciencedirect.com/science/article/pii/S0042698915001649.

Radonjic, A., Cottaris, N. P., & Brainard, D. H. (2015a). Color Constancy in a naturalistic, goal-directed task. Journal of Vision, 15(11), 1–21.

Radonjic, A., Cottaris, N. P., & Brainard, D. H. (2015b). Color constancy supports crossillumination color selection. Journal of Vision, 15(6), 1–19. http://journalofvision.org/content/15/6/13

Radonjic, A., Cottaris, N. P., & Brainard, D. H. (2018). Quantifying how humans trade off color and material in object identification. Paper presented at the Electronic Imagining 2018. Retrieved.

Ramsay, J. O. (1977). Maximum likelihood estimation in multidimensional scaling. Psychometrika, 42, 241–266.

Rodieck, R. W. (1998). The First Steps in Seeing. Sunderland, Mass.: Sinauer.

Rogers, M., Knoblauch, K., & Franklin, A. (2016). Maximum likelihood conjoint measurement of lightness and chroma. Journal of the Optical Society of America A, 33(3), A184–A193. http://josaa.osa.org/abstract.cfm?URI=josaa-33-3-A184.

Saarela, T., & Olkkonen, M. (2017). Integration of color and gloss in surface material discrimination. Journal of Vision, 17(10), 229–229. http://dx.doi.org/10.1167/17.10.229.

Shepard, R. N., & Cooper, L. A. (1992). Representation of Colors in the Blind, Color-Blind, and Normally Sighted. Psychological Science, 3(2), 97–104. https://doi.org/10.1111/j.1467-9280.1992.tb00006.x.

Stockman, A., & Sharpe, L. T. (2000). Spectral sensitivities of the middle-and long-wavelength sensitive cones derived from measurements in observers of known genotype. Vision Research, 40, 1711–1737.

Wandell, B. A. (1995). Foundations of Vision. Sunderland, MA: Sinauer.

Watson, A. B. (2017). QUEST+: A general multidimensional Bayesian adaptive psychometric method. Journal of Vision, 17(3), 1–27.

Wenzel, J. (2014). Mitsuba Documentation (Version 0.5.0).

